# A comparison of pedigree, genetic, and genomic estimates of relatedness for informing pairing decisions in two critically endangered birds

**DOI:** 10.1101/721118

**Authors:** Stephanie Galla, Roger Moraga, Liz Brown, Simone Cleland, Marc P. Hoeppner, Richard Maloney, Anne Richardson, Lyndon Slater, Anna W. Santure, Tammy Steeves

**Affiliations:** School of Biological Sciences, University of Canterbury, Christchurch, New Zealand; Tea Break Bioinformatics, Ltd., Palmerston North, New Zealand; New Zealand Department of Conservation, Twizel, New Zealand; Institute for Clinical Molecular Biology, Christian-Albrechts-University Kiel, Kiel, Germany; New Zealand Department of Conservation, Christchurch, New Zealand; The Isaac Conservation and Wildlife Trust, Christchurch, New Zealand; New Zealand Department of Conservation, Rangiora, New Zealand; School of Biological Sciences, University of Auckland, Auckland, New Zealand

**Keywords:** Conservation genetics, conservation genomics, relatedness, conservation breeding, pairing recommendations, PMx.

## Abstract

Conservation management strategies for many highly threatened species include conservation breeding to prevent extinction and enhance recovery. Pairing decisions for these conservation breeding programmes can be informed by pedigree data to minimise relatedness between individuals in an effort to avoid inbreeding, maximise diversity, and maintain evolutionary potential. However, conservation breeding programmes struggle to use this approach when pedigrees are shallow or incomplete. While genetic data (i.e., microsatellites) can be used to estimate relatedness to inform pairing decisions, emerging evidence indicates this approach lacks precision in genetically depauperate species, and more effective estimates will likely be obtained from genomic data (i.e., thousands of genome-wide single nucleotide polymorphisms, or SNPs). Here, we compare relatedness estimates using pedigree-, genetic-, and genomic-based approaches for making pairing decisions in two critically endangered birds endemic to New Zealand: kakī/black stilt (*Himantopus novaezelandiae*) and kākāriki karaka/orange-fronted parakeet (*Cyanoramphus malherbi*). Our findings indicate genetic-based estimates of relatedness are indeed the least precise when assessing known parent-offspring and full sibling relationships. Furthermore, our results show that relatedness estimates and subsequent pairing recommendations using *PMx* are most similar between pedigree- and genomic-based approaches. These combined results indicate that in lieu of robust pedigrees, SNPs are an effective tool for informing pairing decisions, which has exciting implications for many poorly pedigreed conservation breeding programmes worldwide.

## 1. Introduction

In order to recover the world’s rarest species, a multifaceted approach is needed to address the factors that cause species decline and those that promote species resilience (Grueber et al., 2019; Jamieson 2015; Soulé, 1985). A critical facet of threatened species recovery is genetic management (Frankham, 2005; O’Grady et al., 2006; Spielman, Brook, & Frankham, 2004), including conservation breeding, where breeding individuals in intensively managed captive or wild populations are paired to minimise inbreeding and to maximise genetic diversity in an effort to maintain evolutionary potential (Ballou & Lacy, 1995; Ballou et al., 2010; de Villemereuil et al., 2019; Giglio, Ivy, Jones, & Latch, 2016; Ivy & Lacy 2012). In these conservation breeding programmes, offspring may remain in intensively managed captive or wild populations (e.g., the Tasmanian devil, *Sarcophilius harissii*, insurance population, Hogg et al. 2015; kākāpō, *Strigops habroptilus*; Elliott, Merton, & Jansen, 2001; respectively), or they may be translocated to the wild (e.g., California condor, *Gymnogyps californianus*; Walters et al., 2010). In addition to demographic considerations (e.g., Moore, Converse, Folk, Runge, & Nesbitt, 2012; Slotta-Bachmayr, Boegel, Kaczensky, Stauffer, & Walzer, 2004; Tenhumberg, Tyre, Shea, & Possingham, 2004), current best practice for making pairing decisions in conservation breeding programmes is to use available ancestry data from multigenerational pedigrees to estimate kinship - a metric of pairwise coancestry or relatedness - between all living individuals in a population (Ballou & Lacy, 1995; Ballou et al., 2010; Ivy & Lacy, 2012; Lacy, 1995). Individuals are typically paired to avoid inbreeding and individuals with the lowest mean kinship (i.e., average pairwise relatedness among all others in the population, including oneself) are prioritised for pairing to maximise founder representation (Lacy, 2012; Willoughby et al., 2015).

While pedigrees are still considered the ‘gold standard’ for estimating relatedness in conservation breeding programmes (Hammerly, de la Cerda, Bailey, & Johnson, 2016; Jiménez-Mena, Schad, Hanna, & Lacy, 2016), there are inherent assumptions that, when violated, hinder pedigree accuracy. For example, pedigrees assume that all founders are unrelated (Ballou, 1983), which is unlikely for many highly threatened wild populations, given most will have experienced one or more historical population bottlenecks and founders sourced from these remnant wild populations are likely related (Bergner, Jamieson, & Robertson, 2014; Hogg et al., 2018). Simulation studies suggest that complete pedigrees with substantial depth (> 5 generations recorded) are robust enough to reflect true relatedness and inbreeding estimates despite violating this assumption (Balloux, Amos, & Coulson, 2004; J. Pemberton, 2004; Rudnick & Lacy, 2008). However, in many conservation breeding programmes, unknown founders are routinely sourced from wild populations to supplement captive populations (e.g., kākāriki karaka, *Cyanoramphus malherbi*, this manuscript) and to reduce the risk associated with adaptation to captivity (Frankham, 2008). Under these circumstances, pedigree depth may remain perpetually shallow and therefore susceptible to violations of the assumption of founder relatedness. This is captured by Hogg et al. (2018), who demonstrate how violating this assumption led to a significant underestimation of both relatedness and inbreeding in a conservation breeding programme routinely augmented by wild individuals. In addition to these caveats, many intensively managed populations are poorly pedigreed, meaning these pedigrees contain missing information (i.e., unknown parentage due to matings that include unidentified individuals or extra-pair parentage; Bérénos, Ellis, Pilkington, & Pemberton, 2014; Lacy, 2009; Pemberton, 2008; Putnam & Ivy, 2014) or record keeping errors (e.g., Hammerly et al., 2016).

Even when pedigrees are of high depth, have no missing information, and contain no errors, expected relatedness between individuals can differ from realised relatedness, as pedigrees are based on probabilities as opposed to empirical estimates of genome sharing (Hill & Weir, 2012; Kardos, Luikart, & Allendorf, 2015; Speed & Balding, 2015; Willoughby et al., 2015). Based on Mendelian inheritance, pedigrees estimate the probability that two alleles, one chosen at random from each of two individuals, are identical by descent (IBD) from a parent or common ancestor (Ballou, 1983; Lacy, 1995). When using a pedigree, the relatedness coefficient (*R*) for parents and offspring, as well as for full siblings, is 0.5 when inbreeding is not present, indicating each pair shares 50% of their genomic information. While parents do contribute roughly 50% of their genomic information to their gametes, the combined effects of recombination, independent assortment, and random fertilisation can lead to a larger range of realised relatedness between full siblings (Hill & Weir, 2011, 2012; Speed & Balding, 2015). For example, a simulation study in humans revealed that realised relatedness between full siblings could range anywhere from 0.37-0.61 (Visscher et al., 2006), however this range can vary depending on the genome architecture of the species in question (e.g., number and size of chromosomes and the frequency and location of recombination events; Hill & Weir, 2011; Kardos et al., 2015; Ulrich Knief, Kempenaers, & Forstmeier, 2017).

An alternative approach for populations lacking robust pedigrees is to use genetic-based estimates of pairwise relatedness to inform pairing decisions (Attard et al., 2016; Pemberton, 2004; Pemberton, 2008; Slate et al., 2004). This approach typically uses 8-30 microsatellite markers and empirical allele frequencies to estimate the probability that shared alleles are IBD from a common ancestor (Speed & Balding, 2015). To date, numerous conservation programmes have used this approach to inform pairing recommendations, repair studbooks, and resolve unknown parentage assignments, including programmes for the near-threatened parma wallaby (*Macropus parma*; Ivy, Miller, Lacy, & DeWoody, 2009), the vulnerable Jamaican yellow boa (*Epicrates subflavus*; Tzika, Remy, Gibson, & Milinkovitch, 2009), the critically endangered Anegada iguana (*Cyclura pinguis*; Mitchell et al., 2011), and the critically endangered Attwater’s prairie-chicken (*Tympanuchus cupido attwateri*; Hammerly et al., 2016; Hammerly, Morrow, & Johnson, 2013). While some empirical research indicates that a diverse panel of microsatellites produces diversity estimates that are representative of genome-wide diversity and can be more useful than shallow pedigrees (e.g., Forstmeier, Schielzeth, Mueller, Ellegren, & Kempenaers, 2012), more recent simulation studies indicate that microsatellites provide a poor-indicator of pairwise relatedness and inbreeding, particularly in genetically depauperate endangered species where allelic diversity is low (i.e., < 4 alleles per locus in the founding population; Robinson, Simmons, & Kennington, 2013; Taylor, 2015; Taylor, Kardos, Ramstad, & Allendorf, 2015). These studies argue that a better indication of genome-wide diversity can be obtained from genomic-based estimates of relatedness based on large numbers of genome-wide single nucleotide polymorphisms (i.e., SNPs; Knief et al., 2015; Taylor, 2015; Taylor et al., 2015).

Given the decreasing cost of high-throughput sequencing (Hayden, 2014) and the increasing amount of genomic resources readily available for non-model species (Galla et al., 2019), producing thousands of SNPs is now possible for many highly threatened species and provides an exciting opportunity for use in conservation breeding programmes (Galla et al., 2016; He, Johansson, & Heath, 2016). Indeed, there are several recent examples of genome-wide SNPs being used for relatedness in conservation, ecology, and evolution (e.g., De Fraga, Lima, Magnusson, Ferrão, & Stow, 2017; Escoda, González-Esteban, Gómez, & Castresana, 2017), with some studies indicating that genome-wide SNPs provide greater accuracy in estimating relatedness and inbreeding compared to robust pedigrees (Kardos et al., 2015; Santure et al., 2010; Wang, 2016) or microsatellites (Attard, Beheregaray, & Möller, 2018; Bérénos, Ellis, Pilkington, & Pemberton, 2014; Hellmann et al., 2016; Keller, Visscher, & Goddard, 2011; Lemopoulos et al., 2019; Li, Strandén, Tiirikka, Sevón-Aimonen, & Kantanen, 2011).

To our knowledge, no study has compared pedigree-, genetic-, and genomic-based approaches for estimating relatedness to inform pairing decisions for conservation breeding programmes, despite there being over 350 vertebrates worldwide that are captive bred for release to the wild (Smith et al., 2011). Here, we evaluate the effectiveness of these three approaches using two critically endangered birds endemic birds to Aotearoa New Zealand — the kakī/black stilt (*Himantopus novaezelandiae*) and kākāriki karaka/orange-fronted parakeet (*Cyanoramphus malherbi*) — as Proof-of-Concept. Kakī and kākāriki karaka are excellent candidates for this research as both have active conservation breeding programmes, as well as available multigenerational pedigrees, microsatellites panels (Andrews, Hale, & Steeves, 2013; Steeves, Hale, & Gemmell, 2008) and genomic resources including species-specific reference genomes and whole genome resequencing data (Galla et al., 2019; this study).

Once found on both main islands of Aotearoa, kakī experienced significant population declines throughout the 20th century due to introduced mammalian predators (e.g., cats, stoats, and hedgehogs) along with braided river habitat loss and degradation (Sanders & Maloney, 2002). Today, an estimated 129 kakī are largely restricted to braided rivers of Te Manahuna/The Mackenzie Basin (Department of Conservation, *personal comm.*; Figure 1A) and recovery efforts include a conservation breeding programme that was initiated in the early 1980’s (Reed, 1998). In addition to breeding birds in captivity, the kakī recovery programme also collects eggs from intensively monitored wild nests and rears them in captivity before wild release. Similar to kakī, kākāriki karaka were also once found on both main islands of New Zealand and experienced population declines in the 19th and 20th centuries due to introduced mammalian predators (e.g., brush-tailed possums and stoats) and habitat loss (Kearvell & Legault, 2017). Today, breeding populations of an estimated 100-300 kākāriki karaka are restricted to beech forests in three North Canterbury Valleys (the Hawdon, Hurunui, and Poulter) and to Oruawairua/Blumine Island in the Marlborough Sounds (Department of Conservation, *unpublished data*; Figure 1B). Recovery efforts include a conservation breeding programme initiated in 2003, with founders sourced from the Poulter, Hawdon, and Hurunui Valleys. In most instances, offspring from pairings are released to the Hurunui Valley for wild supplementation. More recently, offspring are also released into the Poulter Valley to encourage pairing with the remaining birds from an extremely small remnant wild population (Department of Conservation, *personal comm.*). Eggs from these pairs are harvested, brought into captivity, and fostered under surrogate birds, with hatchlings incorporated into the conservation breeding programme.

**Figure 1.**
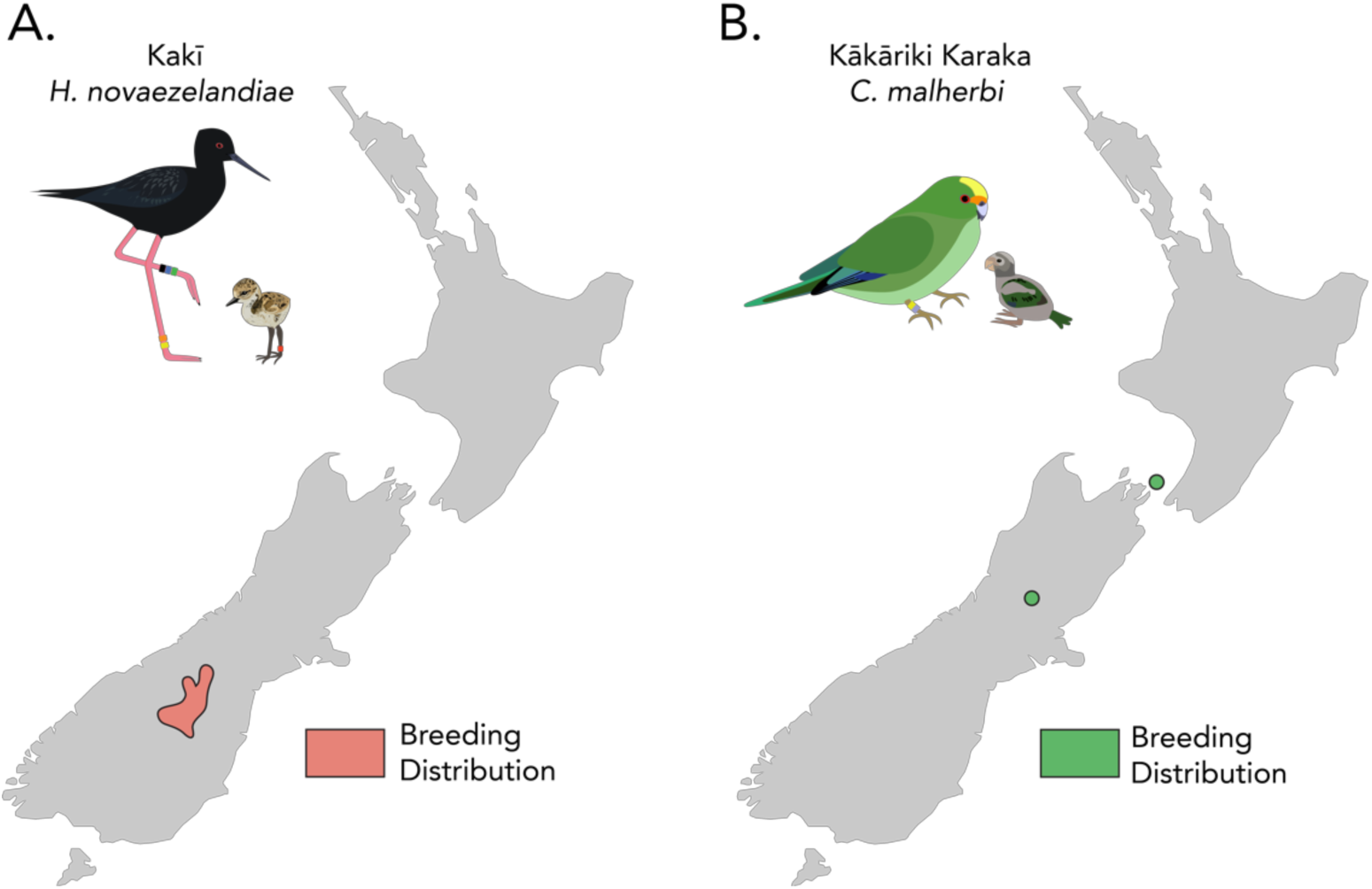
Current breeding distributions of wild kakī (A) and kākāriki karaka (B) in Aotearoa.

Because captive breeding pairs for both species are kept in separate enclosures with offspring intensively managed, kakī and kākāriki karaka present an opportunity to examine relatedness in known family groups. Specifically, we compare relatedness estimates from pedigree, microsatellites, and genome-wide SNPs using known parent-offspring and full sibling relationships. We then compare pairing recommendations among these three approaches to assess how each translates to effective conservation management. While kakī and kākāriki karaka have comparative data sets available, they represent two taxonomically distinct bird species with different life history strategies. Thus, we anticipate the results of our research may be applicable to the wider conservation breeding community.

## 2. Materials and methods

### 2.1. Sample collection and DNA extraction

Animal ethics approval for this project has been granted by the New Zealand Department of Conservation (i.e., DOC; AEC 283). Captive kakī and kākāriki karaka are managed by DOC at two facilities in Aotearoa: the DOC Kakī Management Centre in Twizel and Isaac Conservation and Wildlife Trust in Christchurch. Kakī used in this study are 36 individuals sampled between 2014-2017, including 24 individuals from six captive family groups and 12 individuals from wild parents that represent diverse lineages based on the pedigree. Kākāriki karaka sampled in this study are 36 individuals sampled between 2015-2019, including individuals from eight captive family groups and one wild individual from the Poulter Valley of North Canterbury (Table 1).

**Table 1.**
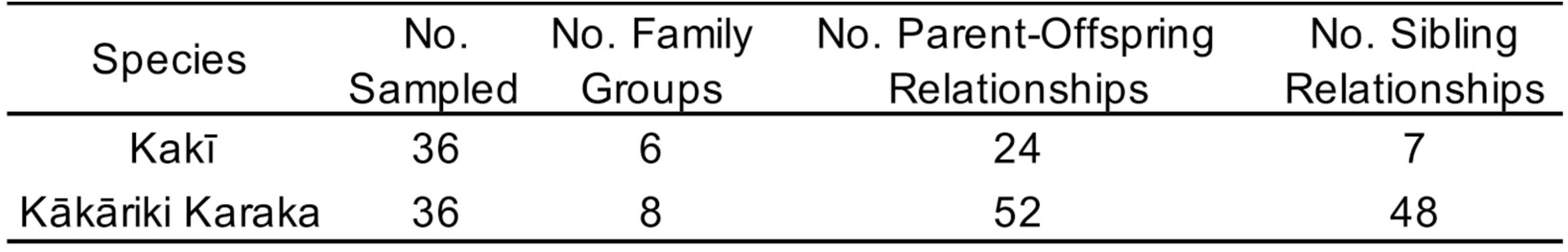
Family group sampling strategy used in this study.

| Species | No. Sampled | No. Family Groups | No. Parent-Offspring Relationships | No. Sibling Relationships |
| --- | --- | --- | --- | --- |
| Kakī | 36 | 6 | 24 | 7 |
| Kākāriki Karaka | 36 | 8 | 52 | 48 |

Blood, feather, or tissue samples were sampled from each bird during routine health checks by DOC and Isaac Conservation and Wildlife Trust staff and immediately transferred into 95% molecular grade ethanol and stored at −80°C. High quantity and quality DNA was extracted using a lithium chloride chloroform extraction method (Galla et al., 2019) at the University of Canterbury School of Biological Sciences. Extractions were assessed for quality by running 2µL of DNA on a 2% agarose gel. A Qubit® 2.0 Fluorometer (Fisher Scientific) was used for DNA quantification.

Known parent-offspring relationships were verified through an allele mismatch exclusion analysis (Jones & Ardren, 2003) using microsatellite panels previously developed for kakī (Steeves et al., 2008) and kākāriki karaka (Andrews, 2013), with a maximum allowed mismatch of one allele at one locus (see *Microsatellite data* below).

### 2.2. Pedigree-based Relatedness

Multigenerational pedigrees were constructed for kakī and kākāriki karaka by entering studbook information (i.e., hatch date, sex, parentage, and status) into the programme PopLink v. 2.5.1 (Faust, Bergström, Thompson, & Bier, 2018). Sex for all individuals was verified using molecular markers 2550F/2718R (Fridolfsson & Ellegren, 1999) for kakī and P2/P8 (Griffiths, Double, Orr, & Dawson, 1998) for kākāriki karaka, with PCR products run on a 3% agarose gel for visual characterisation, with positive controls included. Due to the short distance between P2/P8 alleles on the Z and W chromosomes in kākāriki karaka (Robertson & Gemmell, 2006), 2µL of PCR products using a tagged forward primer were combined with 11.7µL formamide and 0.3µL of Genescan™ LIZ® 500 size standard (Applied Biosystems) and genotyped on an ABI 3739xl (Applied Biosystems), with alleles manually scored using GeneMarker v. 2.2 (State College, PA, USA). Inconsistencies in pedigrees were identified using the validation tool in PopLink and corrected using observations by DOC and the Isaac Conservation and Wildlife Trust. Pedigree depth for kakī is deeper (1-8 generations, 3.35 average) than for kākāriki karaka (1-5 generations, 2.48 average). Pairwise estimates of kinship and inbreeding were produced using the programme PMx v. 1.6.20190628 (Lacy, 2012). Pairwise coefficients of relatedness (*R*) were calculated from kinship data using *R*(*xy*) *=* 2* *f*(*xy*) */ √*{(1+*Fx*)(1+*Fy*)}. In this formula, *f*(*xy*) is the kinship between two individuals (*x* and *y*) and *Fx* and *Fy* are the inbreeding coefficients of individuals *x* and *y* (Crow & Kimura, 1970).

### 2.3. Genetic and genomic-based relatedness

#### 2.3.1. Microsatellite data

Microsatellite loci (*n* = 8) previously described for kakī were amplified using PCR protocols by Steeves *et al.* (Steeves et al., 2008). Microsatellite loci (*n* = 17) designed for kākāriki karaka and one locus (*Cfor0809*) for Forbe’s parakeet (*C. forbesi*) that cross-amplified in kākāriki karaka were amplified using PCR protocols by Andrews *et al.* (2013) and Chan *et al.* (2005), respectively. Samples were prepared for genotyping by adding 0.5 µl of PCR product to 11.7µl formamide and 0.3µl Genescan™ LIZ® 500 size standard (Applied Biosystems) and were genotyped on either a 3130xl or 3730xl Genetic Analyser (Applied Biosystems). Chromatograms were visualised using GeneMarker v. 2.2 (State College, PA, USA). To avoid bias by potential dye shifts (Sutton, Robertson, & Jamieson, 2011), peaks were scored manually. The number of alleles and standard estimates of per locus diversity — including expected heterozygosity (*H_O_*) and observed heterozygosity (*H_E_*) — were produced using GenAlEx v. 6.5 (Peakall & Smouse, 2006; Smouse & Peakall, 2012). Tests for deviations from Hardy-Weinberg and linkage disequilibrium for these loci using samples that are representative of larger kakī and kākāriki karaka populations can be found in Steeves *et al*. (2008; 2010) and Andrews (2013) respectively. For kākāriki karaka, only eight of the 18 microsatellite markers previously described were polymorphic in this study and these eight loci were used in all downstream analyses.

Genetic-based *R* estimates were produced in the programme COANCESTRY v. 1.0.1.9 (Wang, 2011). COANCESTRY offers seven different estimators of relatedness, and to choose the most appropriate estimator for the kakī and kākāriki karaka microsatellite data sets, we employed the simulation module within COANCESTRY using allele frequencies, missing data, and error rates from our microsatellite data sets. To produce dyads that represent the relationships and degree of inbreeding found within kakī and kākāriki karaka, we used R package ‘identity’ (Li, 2010) to generate 10,879 dyads for kakī and 1,484 dyads for kākāriki karaka based on the pedigrees of both species. The frequency of each unique dyad in the kakī and kākāriki karaka data sets were scaled to create 1,000 dyads for each set that are representative of relationships between individuals used in this study. The COANCESTRY simulations were conducted using allele frequencies, error rates, and missing data rates from each microsatellite data set, with settings changed to account for inbreeding. The triadic likelihood approach (Wang, 2007) was selected given it had the highest Pearson’s correlation with ‘true’ relatedness for both data sets (see Supplemental Materials for details). This approach is also preferred, as it is one of the few estimators that accounts for instances of inbreeding (Wang, 2007).

To estimate *R* with our genetic data set, COANCESTRY programme parameters were set to account for inbreeding, with the number of reference individuals and bootstrapping samples set to 100.

#### 2.3.2. Genomic data

To produce genome-wide SNPs for this study, a reference-guided low coverage resequencing approach was used for kakī and kākāriki karaka.

##### 2.3.2.1. Reference genomes

A reference genome for kakī has already been assembled (Galla et al., 2019) and was used in this study. To assemble a *de novo* reference genome for kākāriki karaka, a paired-end library was prepared at the Institute of Clinical Molecular Biology (IKMB) at Kiel University using the Nextera™DNA Flex Library Prep Kit according to manufacturer specifications and sequenced on an Illumina NovaSeq™6000 with 2 x 150 bp reads at a depth of approximately 70x.

FastQC v. 0.11.8 (Andrews, 2010) was used to evaluate the quality of the raw Illumina data and assess potential sample contamination. Initial read trimming was performed using TrimGalore v. 0.6.2 (Krueger, 2019) and Cutadapt v. 2.1 (Martin, 2011) with a median Phred score of 20, end trim quality of 30, a minimum length of 54, and using the --nextseq two-color chemistry option. Kmer analyses were performed using Jellyfish v. 2.2.10 (Marçais & Kingsford, 2011) prior to assembly to assess heterozygosity and contamination. Two genome assembly programmes were tested for assembly performance: Meraculous-2D v. 2.2.5.1 (Chapman et al., 2011) and MaSuRCA v. 3.2.9 (Zimin et al., 2013). Meraculous was run using trimmed reads in diploid mode 1, with all other assembly parameters set to default. MaSuRCA was run using untrimmed reads, as it incorporates its own error correction pipeline. MaSuRCA parameters adjustments include a grid batch size of 300,000,000, the longest read coverage of 30, a Jellyfish hash size of 14,000,000,000, and the inclusion of scaffold gap closing; all other parameters were set to default for non-bacterial Illumina assemblies. The final assembly using the Meraculous pipeline was more fragmented (i.e., an N50 of 28.5 kb with 67,046 scaffolds > 1 kb), while the MaSuRCA genome was less fragmented (i.e., an N50 of 107.4 kb with 66,212 scaffolds > 1 kb) but contained possible artefacts due to heterozygosity (i.e., tandem repeats flanking short stretches of “N”s). To correct for these issues, the Meraculous assembly was first aligned to the MaSuRCA assembly using Last v. 959 (Kielbasa, Wan, Sato, Horton, & Frith, 2011), then alignments were filtered to find matches where the Meraculous assembly spans the entirety of the tandem repeat in the MaSuRCA scaffolds, but lacking the tandem repeat or stretch of “N”s (i.e., gaps). In those cases, the aligned sequence in the MaSuRCA scaffold was replaced with the Meraculous match.

##### 2.3.2.2. Whole-genome resequencing

Kakī resequencing libraries were prepared at IKMB using a TruSeq® Nano DNA Library Prep kit following the manufacturer’s protocol and were sequenced across 34 lanes of an Illumina HiSeq 4000. 24 individuals were sequenced at high coverage depth (approximately 50x) for an aligned study, and all others were sequenced at a lower coverage depth (approximately 10x). Kākāriki karaka libraries were prepared at IKMB using the Nextera™DNA Flex Library Prep Kit according to manufacturer specifications and sequenced across one lane of an Illumina NovaSeq™6000 at IKMB at a coverage depth of approximately 10x, with one individual sequenced at a depth of 70x, which was additionally used for the reference genome (see above).

FastQC v. 0.11.4 and 0.11.8 (S. Andrews, 2010) were used to evaluate the quality of the raw Illumina data for kakī and kākāriki karaka, respectively. Kakī resequencing reads were subsequently trimmed for the Illumina barcode, a minimum Phred quality score of 20, and a minimum length of 50bp using Trimmomatic v. 0.38 (Bolger, Lohse, & Usadel, 2014). Because kākāriki karaka libraries were produced using different library preparation protocols and nextera chemistry, reads were trimmed using TrimGalore v. 0.6.2 (Krueger, 2019) for nextera barcodes and two-colour chemistry, using a median Phred score of 20, end trim quality of 30, and a minimum length of 54. Prior to mapping, the kakī reference genome was concatenated to a single chromosome using the custom perl script ‘concatenate_genome.pl’ (Moraga, 2018a) for use in an aligned project that used both resequencing and genotyping-by-sequencing reads (see Galla et al., 2019). The kakī and kākāriki karaka reference genomes were indexed and resequencing reads were mapped using Bowtie2 v. 2.2.6 and v. 2.3.4.1 (Langmead & Salzberg, 2012), respectively, with the setting *--very-sensitive*. Resulting SAM files were converted to BAM and were sorted using Samtools v. 1.9. (Li et al., 2009). Read coverage and variant calling were performed using *mpileup* in BCFtools v. 1.9 (Li et al., 2009). The custom perl script ‘split_bamfile_tasks.pl’ (Moraga, 2018b) was used to reduce the computational time needed for mpileup by increasing parallelisation. SNPs were detected, filtered, and reported using BCFtools. Filtering settings were set to retain biallelic SNPs with a minor allele frequency (MAF) greater than 0.05, a quality score greater than 20, and a maximum of 10% missing data per site. After a series of filtering trials for each species (see Supplemental Materials for details), depth for kakī was set to have an average mean depth greater than 10, while kākāriki karaka depth was set so that each site had a minimum depth of 5 and a maximum depth of 200. Resulting SNPs for both data sets were pruned for linkage disequilibrium using BCFtools with the *r^2^* set to 0.6 and a window size of 1000 sites. Sites were not filtered for Hardy-Weinberg equilibrium, as the nature of these data sets (mostly family groups) violates the assumptions of random mating. Per site missingness, depth, and diversity — including proportion of observed and expected heterozygous SNP sites per individual, nucleotide diversity, and SNP density per kb — were evaluated in the final sets using VCFtools v. 1.9 (Danecek et al., 2011).

##### 2.3.2.3. SNP-based relatedness

To produce estimates of *R* using whole-genome SNPs, the programme KGD (Dodds et al., 2015) was used, as it was designed to estimate relatedness using reduced-representation and resequencing data while taking into account read depth. Furthermore, KGD approximates parent-offspring relatedness estimates closer to 0.5 compared to the triadic likelihood approach method (Wang, 2007), which underestimates relatedness, while still providing estimates that significantly correlate with this traditional estimator in both kakī (Pearson’s *r* = 0.96, *p* < 0.001) and kākāriki karaka (Pearson’s *r* = 0.96, *p* < 0.001; see Supplemental Materials for details). Pairwise *R* values derived from KGD were scaled so that self-relatedness for all individuals was equal to 1 using the formula *MS = D x MO x D* where *MS* is the scaled matrix, *MO* is the original matrix, and *D* is a diagonal matrix with elements *D = 1/√(diag(MO))*.

### 2.4. Comparison of relatedness

Pedigree, microsatellite, and SNP-based *R* between all individuals (*n* = 630) were compared among the three methods using Pearson’s correlation coefficient (*r*). While our relatedness data sets are non-parametric, Pearson’s was used over non-parametric tests, such as rank correlations, as our pedigree and microsatellite data sets have an excess of tied values.

### 2.5. Pairing recommendations

We used mate suitability index (MSI) scores in PMx v. 1.6.20190628 (Lacy, 2012) to show how pairing recommendations may change using pedigree-, microsatellite-, and SNP-based approaches for estimating *R*. MSI scores indicate how valuable offspring of a potential pair would be to the population on a scale from 1-6, with 1 being “very beneficial”, and 6 being “very detrimental”. An additional category, usually denoted with a “-” indicates that a pairing is “very highly detrimental” based on a high degree of kinship, and therefore the pairing should be avoided; for these analyses, this category has been put on the numerical MSI scale as a 7. MSI score takes into account four factors: deltaGD (i.e., the net positive or negative genetic diversity provided to the population), the difference of mean kinship values of the pair, the inbreeding coefficients of resulting offspring, and unknown ancestry (Ballou et al. 2001; Ballou et al. 2011). Pairing recommendations using the pedigree were made treating all individuals with unknown parentage in the pedigree as founders. To produce pairing recommendations using genetic and genomic markers, empirical *R* estimates were uploaded to PMx and weighted to 1 to produce MSI scores relying only on empirical data.

Pearson’s correlation (*r*) was used to evaluate whether pairwise MSI scores between the approaches were concordant. To test whether the distribution of MSI scores were statistically different from one another, we used a non-parametric Kruskal-Wallis test with Bonferroni correction and a Tukey Honest Significant Difference (HSD) test.

## 3. Results

### 3.1. Pedigree-based relatedness

This study has produced the first functional multigenerational pedigrees for two critically endangered endemic birds from Aotearoa. The kakī pedigree includes 2,680 wild and captive individuals recorded from 1977-present. Pedigree-based *R* between all kakī used in this study ranged from 0 to 0.56, with an average *R* of 0.13 ± SD 0.13. The average coefficient of relatedness between all known kakī parent-offspring was higher than the expected 0.5 contribution from each parent (0.52 ± SD 0.02), with averaged full sibling *R* of 0.52 ± SD 0.02 (Figure 2). The kākāriki karaka pedigree includes 624 captive individuals from 2003-present. Pedigree-based *R* for all individuals in the pedigree ranged from 0 to 0.67, with an average *R* of 0.19 ± SD 0.18. Average *R* between all parent-offspring was 0.52 ± SD 0.03, with averaged full sibling *R* being 0.51 ± SD 0.02 (Figure 2).

**Figure 2.**
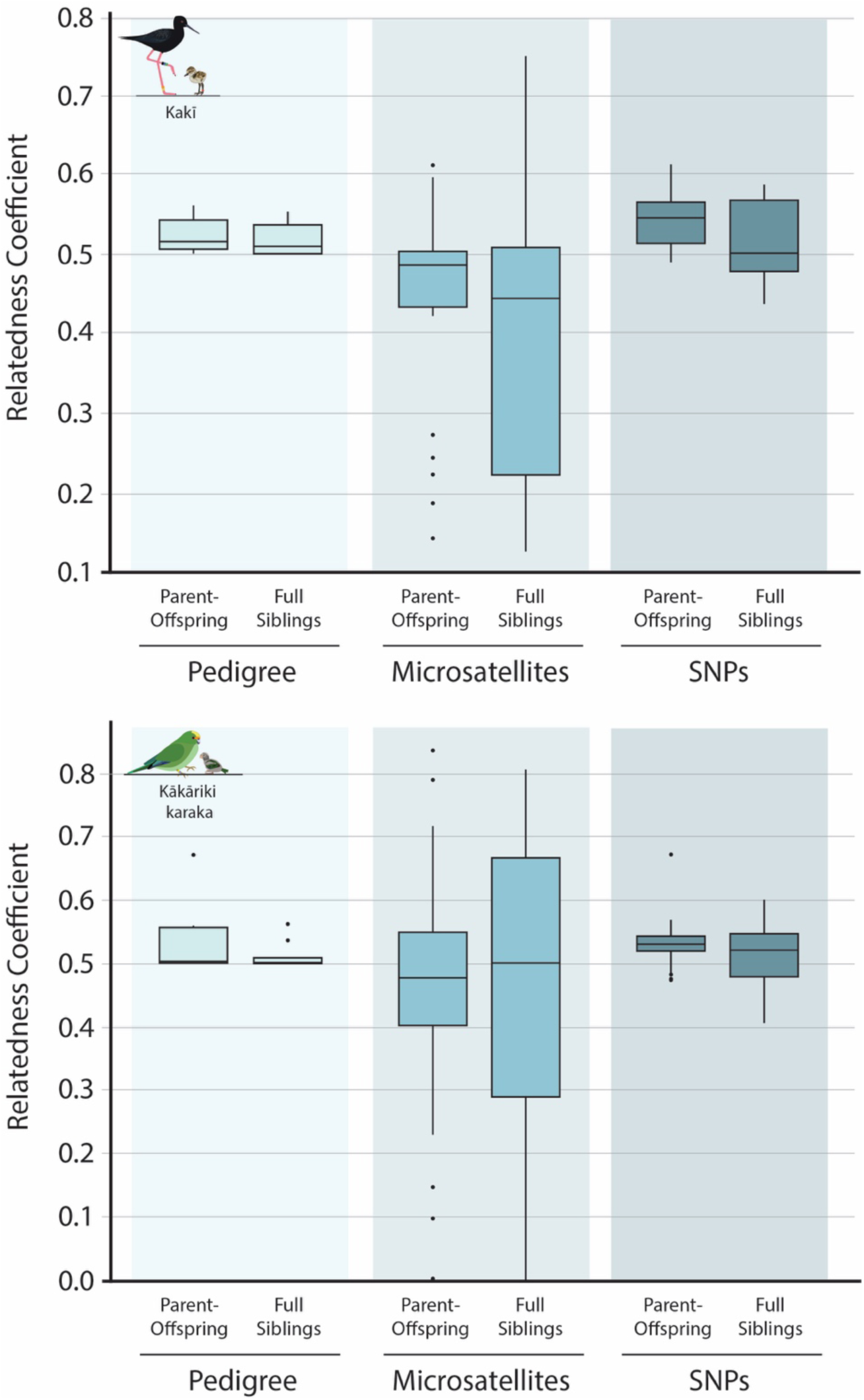
Parent-offspring and full sibling relatedness values derived from pedigree-(pale blue), microsatellite-(medium blue), and SNP-based (dark blue) methods in kakī (top graph) and kākāriki karaka (bottom graph).

### 3.2. Microsatellite diversity and relatedness

All eight microsatellite markers for kakī successfully amplified in all individuals used in this study. The number of alleles present across kakī loci ranged from 2-4 (average 3.13 ± SD 0.64; Table 2), with overall fewer alleles found here than reported in previous studies utilising these loci with more individuals (Hagen, Hale, Maloney, & Steeves, 2011; Steeves et al., 2010). While eighteen microsatellite markers were amplified in kākāriki karaka, one was removed from this study for not successfully amplifying in more than 50% of individuals (locus *OFK56*) and nine were removed for being monomorphic (Table 2). The number of alleles among polymorphic loci ranged from 2-4 (average 3.0 ± SD 0.93), with overall fewer alleles found here than reported in previous studies (Andrews, 2013; Andrews et al., 2013). Observed (*H_O_*) and expected (*H_E_*) heterozygosity for kakī (average *H_O_* = 0.57 ± SD 0.17, average *H_E_* = 0.54 ± SD 0.14) was higher than kākāriki karaka (average *H_O_* = 0.43 ± SD 0.23, average *H_E_* = 0.43 ± SD 0.20; Table 2).

**Table 2.** Descriptive statistics, including number of alleles, observed heterozygosity (H_O_), and expected heterozygosity (H_E_) for microsatellite loci used in this study. Loci from kākāriki karaka that were monomorphic (OFK12, OFK 19, OFK21, OFK26, OFK31, OFK33, OFK52, OFK56, OFK58, OFK61) are not included.

| Species | Locus | No. Alleles | $H_O$ | $H_E$ |
| --- | --- | --- | --- | --- |
| Kakī | BS2 | 3 | 0.667 | 0.652 |
|  | BS9 | 3 | 0.611 | 0.59 |
|  | BS12 | 3 | 0.278 | 0.245 |
|  | BS13 | 2 | 0.528 | 0.5 |
|  | BS21 | 4 | 0.833 | 0.703 |
|  | BS27 | 4 | 0.667 | 0.551 |
|  | BS40 | 3 | 0.444 | 0.448 |
|  | BSdi7 | 3 | 0.556 | 0.596 |
| Kākāriki Karaka | OFK9 | 4 | 0.472 | 0.477 |
|  | OFK41 | 4 | 0.75 | 0.702 |
|  | OFK50 | 3 | 0.444 | 0.513 |
|  | OFK54 | 4 | 0.722 | 0.574 |
|  | OFK55 | 2 | 0.278 | 0.346 |
|  | OFK60 | 2 | 0.222 | 0.239 |
|  | OFK62 | 2 | 0.083 | 0.08 |
|  | C for 809 | 3 | 0.44 | 0.52 |

Microsatellite-based *R* between all kakī used in this study ranged from 0 to 0.85, with an average *R* of 0.16 ± SD 0.19. Average *R* between all known kakī parent-offspring (0.44 ± SD 0.13) was below the expected relatedness value of 0.5, with a larger standard deviation of *R* values compared to pedigree-based estimates. Averaged full sibling *R* (0.41 ± SD 0.20) also had a larger deviation around the mean compared to the microsatellite-based parent-offspring estimates (Figure 2).

Microsatellite-based *R* between all kākāriki karaka used in this study ranged from 0 to 0.84, with an average *R* of 0.18 ± SD 0.22. Similar to kakī, average *R* between all known kākāriki karaka parent-offspring relationships (0.47 ± SD 0.19) was below the expected *R* value of 0.5, with a larger standard deviation of *R* values compared to pedigree-based estimates. Averaged full sibling *R* (0.49 ± SD 0.21) also had a larger deviation around the mean compared to microsatellite-based parent-offspring estimates (Figure 2).

### 3.3. Reference genome assembly, SNP discovery, diversity, and relatedness estimates

#### 3.3.1. Kākāriki karaka reference genome assembly

Reference genome library preparation and Illumina NovaSeq™sequencing resulted in 584.47 million total reads for the kākāriki karaka genome. The final kākāriki karaka genome assembly was 1.15GB in length, which is within the range of many assembled avian genomes (Zhang et al., 2014). The final assembly had 66,212 scaffolds with a scaffold N50 of 107.4 kb. See Data Availability section for access information.

#### 3.3.2. SNP discovery and diversity

Library preparation and Illumina sequencing resulted in 6.07 billion total reads for kakī (168.69 ± SD 65.32 million reads). In addition to the individual used for the reference assembly, 3.64 billion total reads (average = 103.92 ± SD 29.76 million reads) were produced for the additional 35 kākāriki karaka in this study. More SNPs were discovered during initial SNP discovery using kākāriki karaka than kakī, and more remained post filtering (Table 3). These filtered SNPs were used for all downstream analyses. Average missingness was low for both data sets (Table 3), but lower for kākāriki karaka than kakī, as kākāriki karaka had a hard minimum cut-off for depth during filtering that resulted in no missing data. Average depth for both data sets was relatively high (Table 3), with kakī having slightly higher average depth. Average diversity statistics (nucleotide diversity, and the average observed and expected SNP heterozygosity per individual post filtering) were similar in both species, with diversity in kakī being slightly higher. SNP density using the kakī data set was higher than the kākāriki karaka data set, indicating that discovered SNPs are closer in proximity in the kakī data set (Table 3).

**Table 3.** Descriptive statistics, including number of SNPs pre- and post-filtering, average depth, average missingness, average nucleotide diversity (π) ± SD, average proportion of observed heterozygous SNP sites (H_O_) ± SD, average proportion of expected heterozygous SNP sites (H_E_) ± SD, and average SNP density (number of SNPs per kilobase) ± SD.

| Species | No. of SNPs<br>Pre-Filtering | No. of SNPs<br>Post-Filtering | Average<br>Depth | Average<br>Missingness | Average $\pi$ | Average $H_O$<br>of SNP Sites | Average $H_E$<br>of SNP Sites | SNP<br>Density |
| --- | --- | --- | --- | --- | --- | --- | --- | --- |
| Kakī | 4,246,100 | 68,144 | 28.73 ± 10.29 | 0.002 ± 0.004 | 0.35 ± 0.14 | 0.40 ± 0.02 | 0.35 ± 0.00 | 0.58 ± 3.18 |
| Kākāriki Karaka | 22,435,128 | 90,949 | 25.1 ± 14.87 | 0.00 ± 0.00 | 0.33 ± 0.14 | 0.37 ± 0.02 | 0.33 ± 0.00 | 0.17 ± 0.42 |

#### 3.3.3. SNP-based relatedness

SNP-based *R* between all kakī used in this study ranged from 0.13-0.61, with an average *R* of 0.27 ± SD 0.09. Similar to pedigree-based estimates, average *R* between all known kakī parent-offspring were slightly higher than the expected relatedness value of 0.5 with a small standard deviation relative to microsatellite-based estimates (0.54 ± SD 0.03). Averaged full sibling *R* also had a larger deviation around the mean (0.52 ± SD 0.05) than parent-offspring relationships (Figure 2).

SNP-based *R* between all kākāriki karaka used in this study ranged from 0.08-0.67, with an average *R* of 0.30 ± SD 0.12. Similar to pedigree-based estimates, average *R* between all known kākāriki karaka parent-offspring was slightly above the expected *R* value of 0.5 with a small standard deviation relative to genetic-based estimates (0.53 ± SD 0.03). Averaged full sibling relatedness also had a larger deviation around the mean (0.52 ± SD 0.05) compared to the pedigree-based estimates (Figure 2).

### 3.4. Comparison of relatedness estimates and pairing recommendations

All kakī and kākāriki karaka *R* estimates using pedigree-, microsatellite-, and SNP-based approaches correlated with one another with high statistical significance (Pearson’s Correlation, *p* < 0.001; Table 4, Figure 3). Of all the approaches, the correlation coefficient between pedigree- and SNP-based approaches was markedly higher than between other approaches, indicating that they are the most concordant (Figure 3).

**Figure 3.**
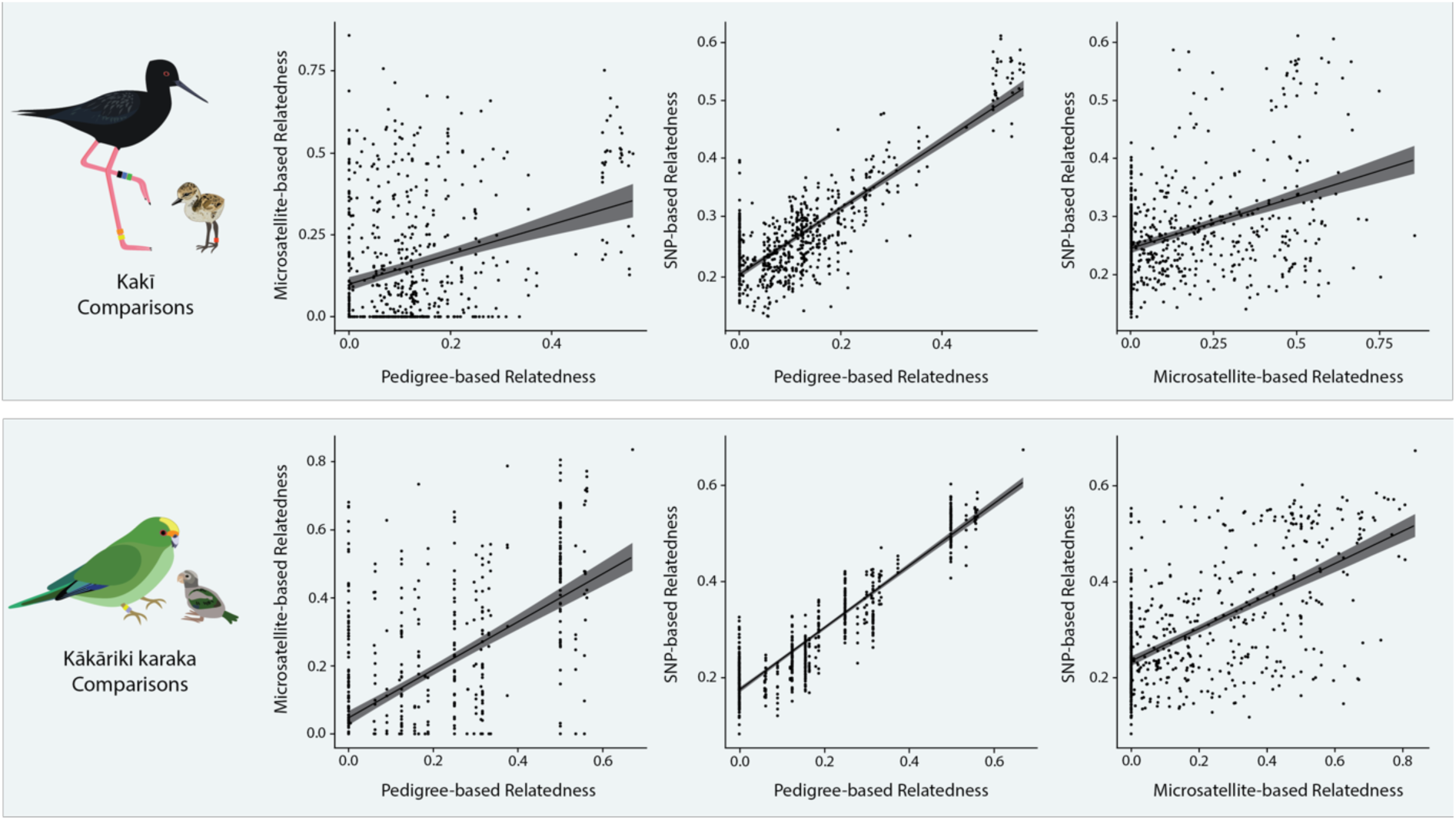
Scatterplots showing relationships between pedigree-, microsatellite-, and SNP-based relatedness estimates in known family groups for kakī and kākāriki karaka. A trend line (black) and 95% confidence intervals (grey) are shown in each comparison.

**Table 4.** Pearson’s correlation coefficients (r) between different relatedness estimators and subsequent MSI scores in kakī and kākāriki karaka. ** equates to statistical significance of p < 0.01, while *** equates to statistical significance of p < 0.001.

| Species | Metric | Pedigree v. Microsatellite | Pedigree v. SNPs | Microsatellite v. SNPs |
| --- | --- | --- | --- | --- |
| Kakī | Relatedness | 0.30*** | 0.83*** | 0.39*** |
|  | MSI Scores | 0.15** | 0.50*** | 0.20*** |
| Kākāriki karaka | Relatedness | 0.57*** | 0.93*** | 0.61*** |
|  | MSI Scores | 0.20*** | 0.65*** | 0.36*** |

The mate-suitability index (MSI) scores were calculated as an approximation for pairing recommendations derived from *R* estimates using the different approaches. Average pedigree-based MSI scores for kakī (4.46 ± SD 1.59) were lower on average than microsatellite-based scores (4.73 ± SD 1.63), but not significantly different from each other (Kruskall-Wallis test with Bonferroni correction, *p* = 0.2). SNP-based MSI scores for kakī (average = 5.67 ± SD 1.39) were significantly higher than pedigree- and microsatellite-based scores (Kruskall-Wallis test with Bonferroni correction, *p* < 0.001), with SNP-based scores providing the highest frequency of category 7 (i.e., highly detrimental) pairings (Figure 4). While the distributions of MSI scores between each approach were different, each approach produced scores that correlated significantly with one another (Pearson’s correlation, *p* < 0.01-0.001). Similar to correlations between *R* estimates, correlation coefficients between pedigree and SNP-based MSI scores were highest (Table 4). Of all possible pairings represented by MSI scores, 38% did not experience a change in score value between pedigree- and SNP-based approaches; however, 20% of pairings experienced an MSI score change that was 3+ categories different. In 2% of pairings, pedigree-based MSI scores were categorised as a 1 (i.e., preferred pairing) while SNP-based MSI scores were categorised as a 7 (i.e., highly detrimental).

**Figure 4.**
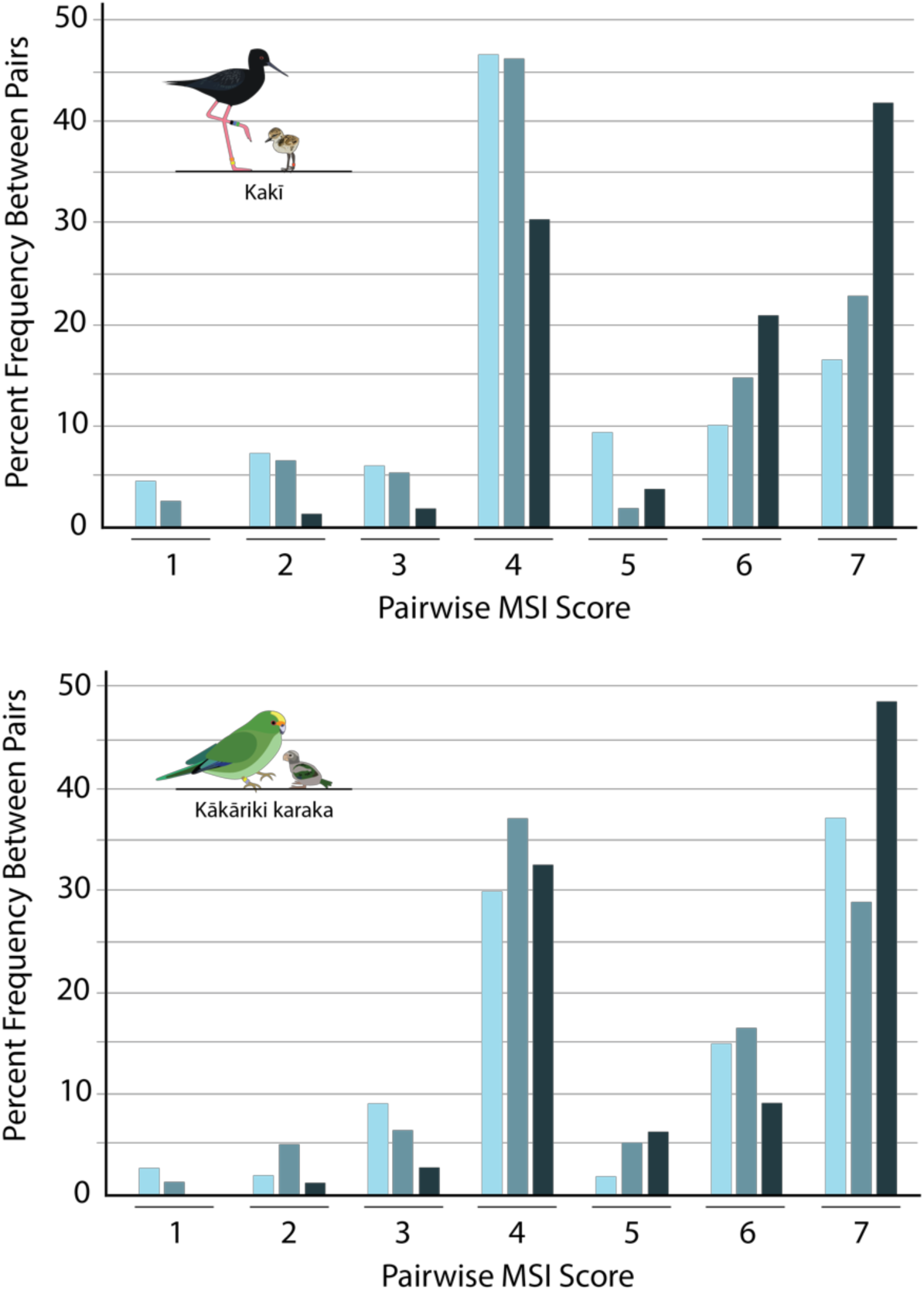
Frequency of MSI scores using pedigree - (pale blue), microsatellite-(medium blue), and SNP-based (dark blue) kinship/relatedness values in kakī and kākāriki karaka.

Similar to kakī, average kākāriki karaka SNP-based MSI scores (5.64 ± SD 1.47) were significantly higher than pedigree (5.20 ± SD 1.71) and microsatellite (5.04 ± SD 1.61) scores (Kruskall-Wallis test with Bonferroni correction, *p* < 0.001), while pedigree- and microsatellite-based scores did not significantly differ (Kruskall-Wallis test with Bonferroni correction, *p* = 0.67). SNP-based scores provided the highest frequency of category 7 (i.e., highly detrimental) pairings (Figure 4). Each approach also produced scores that correlated significantly with one another (Pearson’s correlation, *p* < 0.001), with the highest correlation coefficients seen between pedigree and SNP-based MSI scores (Pearson’s *r* = 0.65; Table 4). Of all possible pairings represented by MSI scores, 59% did not experience a change in score value between pedigree- and SNP-based approaches; however, 9% of pairings experienced an MSI score change that was 3+ categories different. In 2% of pairings, pedigree-based MSI scores were categorised as a 1 (i.e., very beneficial) while SNP-based MSI scores were categorised as a 7 (i.e., highly detrimental).

## 4. Discussion

This study is the first to compare pedigree-, microsatellite-, and SNP-based estimates of relatedness and subsequent pairing recommendations for conservation breeding programmes. The results indicate that microsatellites provide the least precision when estimating relatedness in known family groups, with pedigree- and SNP-based estimates providing higher precision and a much closer approximation of parent-offspring and full sibling relatedness. Further, estimates of relatedness and downstream pairing recommendations are both more similar when using pedigree- and SNP-based data sets compared to microsatellite-based data sets. Despite this, there were important differences in pairing recommendations between the two approaches, with SNP-based mate suitability index (MSI) scores being statistically higher than pedigree-based scores, and some substantial disagreements existing between the two sets of MSI scores. Together, this study provides insight into the differences between pedigree-, microsatellite-, and SNP-based approaches for making pairing recommendations and a pathway for estimating relatedness using genome-wide SNPs to inform pairing decisions in poorly pedigreed conservation breeding programmes worldwide.

### 4.1. Relatedness comparisons

Pedigree-based estimates of parent-offspring and full sibling relatedness approximated 0.5 for both kakī and kākāriki karaka (Figure 2), with some measures being slightly higher, which likely reflects intergenerational inbreeding. These results are consistent with expectations, as pedigrees are based on the probability of Mendelian inheritance, which postulates that first-order relationships (i.e., parents and offspring, and siblings) share 50% of their genomic information (Lacy, 1995; Wright, 1922). We expect realised (i.e., empirical) parent-offspring relationships to also approximate 0.5, but a broader range of realised relatedness estimates among full siblings, as they may receive different genetic material from each parent due to recombination and independent assortment during meiosis, and random fertilisation (Hill & Weir, 2011, 2012; Speed & Balding, 2015). Thus, even when pedigrees are robust, an unavoidable shortcoming is that they do not adequately capture true relatedness between full siblings. We anticipate this uncaptured diversity may prove useful for maximising existing diversity, especially in conservation breeding programmes with relatively few founders (Ballou & Lacy, 1995).

When examining our empirical data sets (i.e., microsatellites and SNPs), more variation is captured between siblings than parents and offspring, albeit with different levels of precision between empirical approaches used (Figure 2). A broad range of microsatellite-based relatedness estimates were observed in both parent-offspring and sibling relationships. In some instances, even parent-offspring pairings appeared relatively unrelated using microsatellites (e.g., minimum parent-offspring *R* = 0.14 in kakī and *R* = 0 in kākāriki karaka), which underscores the lack of precision in this approach and how it could inadvertently lead to poorly informed pairing recommendations. These large ranges of relatedness values using microsatellites can be explained because genetic-based relatedness values between parent-offspring and full siblings are based on allele frequencies, and relatedness between individuals that share common alleles will be substantially lower than individuals that share rare alleles (Speed & Balding, 2015; Wang, 2011). This bias in relatedness values can be exacerbated when samples sizes are small (Wang, 2017), which is typical of conservation breeding programmes. Furthermore, the lack of precision using microsatellites shown here is consistent with studies that suggest relatively few markers with low allelic diversity are insufficient for estimating relatedness and inbreeding, especially in genetically depauperate species (e.g., Attard et al., 2018; Escoda, González-Esteban, Gómez, & Castresana, 2017; Hellmann, Sovic, Gibbs, Reddon, O’Connor, Ligocki, Marsh-Rollo, et al., 2016; Taylor, 2015; Taylor et al., 2015). While the number of microsatellites used here was small and allelic diversity was relatively low, the use of a few microsatellite markers to estimate relatedness and other diversity measures (e.g., inbreeding) is not uncommon (e.g., Hammerly et al., 2013; 2016; Hellmann et al., 2016; Tzika et al., 2009) and low allelic diversity is expected in endangered species (Taylor, 2015; Taylor et al., 2015). Furthermore, our results align with even larger microsatellite panels (e.g., Hoffman et al., 2014; McLennan, Wright, Belov, Hogg, & Grueber, 2019) which show less precision in estimating relatedness and inbreeding than genome-wide SNPs.

Compared to microsatellite-based relatedness, SNP-based relatedness showed a relatively small range with parent-offspring and full sibling relatedness estimates approximating 0.5, and full siblings showing a wider range of values than parent-offspring relationships (Figure 2). Not only is this pattern consistent with expectations given the behaviour of chromosomes during meiosis and random fertilisation, but it also shows more precision than the microsatellite data sets. Other researchers have found similar results in a diverse range of wild taxa, indicating that thousands of genome-wide SNPs show more precision than microsatellites when measuring relatedness and inbreeding (e.g., Attard et al., 2018; Hellmann, Sovic, Gibbs, Reddon, O’Connor, Ligocki, Marsh-Rollo, et al., 2016; Hoffman et al., 2014; Lemopoulos et al., 2019; Thrasher, Butcher, Campagna, Webster, & Lovette, 2018).

Beyond parent-offspring and full sibling relationships, pedigree and SNP-based relatedness estimates showed the highest concordance with one another among the three approaches used (Figure 3). In kakī, the data sets used here include non-captive bred individuals with intensively monitored wild parents. These results provide more credibility to the semi-wild kakī pedigree, where socially monogamous wild pairs of kakī are assumed to be the genetic parents of offspring at nests (but see also Overbeek et al., 2017), as it shows that the kakī pedigree generally concurs with empirical relatedness estimates. Still, it should be noted that many pairs with pedigree-based relatedness values of 0 had pairwise relatedness values ranging upwards of 0.40 in kakī 0.33 and in kākāriki karaka using a SNP-based approach, which approximates first and second order relationships in both species (Figure 2). This indicates that pedigree-based *R* between these individuals may be downwardly biased by unknown founders, missing information, and/or low pedigree depth (Balloux et al., 2004; Bérénos et al., 2014; Hammerly et al., 2016; Hogg et al., 2018; Kardos et al., 2015; Lacy, 1995; Pemberton, 2008; Rudnick & Lacy, 2008; Tzika et al., 2009).

### 4.2. Pairing recommendations

When these relatedness values are translated into pairing recommendations using MSI scores, there is a high concordance between pedigree and SNP-based approaches, despite the SNP-based MSI scores being significantly higher. This latter result is somewhat expected, given that average relatedness estimates using SNPs was highest among the approaches used here, and empirical estimates of relatedness and inbreeding are usually higher than pedigrees as they more effectively capture relatedness between founders or mis-assigned individuals (Hammerly et al., 2016; Hogg et al., 2018). With that said, when making pairing recommendations using kinship-based pairing decisions (e.g., Ballou & Lacy, 1995), it is often the relative kinships between individuals that are more important than absolute values (Galla et al., 2019; McLennan et al., 2019). This suggests that pedigree and SNP-based approaches both yield similar results for pairing recommendations, with some important differences. For example, while correlations between these two sets of MSI scores are high relative to other comparisons, there are still some instances here where pairings are considered ‘highly beneficial’ (i.e., MSI category 1) when using the pedigree and ‘very highly detrimental’ (i.e., MSI category 7) when using SNPs. We expect some differences in MSI scores using pedigree and empirical approaches, given pedigrees MSI scores use kinship values based on Mendelian inheritance while SNP-based estimates are based on the proportion of observed alleles that are identical by state and inferred to be identical by descent given allele frequency information. However, these discrepancies are noteworthy, as they may be indicative of errors in the pedigree (e.g., Hammerly et al., 2016) or affects from founders that are assumed to be unrelated (e.g., Hogg et al. 2018).

### 4.3. Management Implications

Pairing recommendations take into account logistical, demographic, and genetic considerations to maximise population recovery. From these combined results shown here, we recommend that when conservation breeding programmes are poorly pedigreed, SNPs should be used to provide a precise indicator of relatedness to genetically inform pairing decisions. We anticipate SNPs will be particularly applicable in circumstances when pedigrees are the least reliable. For instance, when the founders of a conservation breeding population have no ancestry data available and are likely to be related, SNP-based relatedness estimates between individuals can be used to avoid highly related matings (Hogg et al., 2018). This situation may not only coincide with the original founding event of a captive population, but iteratively when individuals are sourced from wild or translocated populations to augment the captive population, as suggested in Frankham (2008) and Hogg et al (2018). For example, in kākāriki karaka, whole genome resequencing has been made available for all current breeding individuals in the conservation breeding programme, including individuals who are founders themselves. Because birds of unknown ancestry are being routinely sourced from highly endangered wild populations, and will also be founders, we anticipate the need for resequencing these birds as they are incorporated into the breeding programme to assess their relatedness to other individuals.

While we expect SNPs will be important for pairing recommendations moving forward, there are relatively few studies to date that effectively combine existing pedigree data with genomic estimates of relatedness to inform pairing recommendations (but see Hogg et al., 2018; Ivy, Putnam, Navarro, Gurr, & Ryder, 2016). To date, these studies are largely limited to SNPs being used for parentage reconstruction (reviewed in Flanagan & Jones, 2019), where unknown or uncertain relationships are reconstructed using empirical data and software (e.g., Whalen, Gorjanc, & Hickey, 2018), and more complete pedigrees are used moving forward. Alternatively, there is an option to produce empirical estimates of relatedness for all founders or breeding individuals in conservation breeding programmes — as suggested in Ivy et al. (2016) and practiced in Hogg et al. (2018) — and use this baseline of known relatedness moving forward using pedigrees. While the programme PMx allows for the inclusion of empirical data, this approach requires caution, as the calculation of pedigree-based identity by descent for subsequent generations – including kinship and gene diversity — will be affected by the addition of empirical data (Hogg et al. 2018). We acknowledge this approach requires further investigation and validation, particularly for species that receive periodic influx of wild individuals of unknown ancestry in their conservation breeding programme.

### 4.4. Future Directions and Concluding Remarks

Although beyond the scope of this study, we hypothesize that SNPs will provide more useful estimates of relatedness over pedigrees, given they capture genetic variation among full siblings (Hill & Weir, 2011; Kardos et al., 2015; Ulrich Knief et al., 2017; Speed & Balding, 2015). Simulation studies have shown that estimates of identity by descent are more precise when using thousands of genome-wide SNPs than pedigrees (e.g., Kardos et al., 2015; Wang, 2016). Further, a recent study in zebra finch (*Taeniopygia guttata*) by Knief et al. (2017) shows that the amount of Mendelian ‘noise’ resulting from meiotic recombination will be exacerbated in species where crossing over events are non-uniformly distributed, such as birds (Ellegren, 2013). We expect that simulations using pedigree- and SNP-based pairing recommendations over several generations will determine whether a pedigree- or SNP-based approach best maximises genome-wide diversity in the long-term for conservation breeding programmes.

With that said, for poorly pedigreed populations, we recommend a SNP-based approach to estimating relatedness for subsequent pairing recommendations. Given that SNPs have been successfully used to estimate relatedness for different purposes across a wide diversity of taxonomic groups outside of this study (as reviewed in Attard et al., 2018), we anticipate a SNP-based approach for estimating relatedness and making subsequent pairing recommendations will be applicable beyond birds. It should be noted that many approaches used to date have used *de novo* reduced representation approaches (e.g., genotyping-by-sequencing, RADseq; Narum, Buerkle, Davey, Miller, & Hohenlohe, 2013) for SNP discovery, which typically have more missing data, lower depth, and fewer resulting SNPs than the reference-guided whole genome resequencing approach used here. While these factors may contribute to bias in relatedness estimates (but see Dodds et al., 2015), research still indicates that they can provide more precision than microsatellites (Attard et al., 2018). We expect reduced-representation approaches will persist in the short-term, especially for species with massive and complex genomes (e.g., some fish, amphibians, and invertebrates) that otherwise cannot yet be affordably resequenced across entire captive populations. We look forward to seeing more taxonomically diverse species use a SNP-based approach for estimating relatedness, as the results from this paper suggest it can be applied to poorly pedigreed conservation breeding programmes for making pairing recommendations worldwide.

## Acknowledgements

We are grateful for the support of Te Rūnanga o Ngāi Tahu, Te Ngāi Tūāhuriri Rūnanga, Te Rūnanga o Arowhenua, Te Rūnanga o Waihao and Te Rūnanga o Moeraki. We thank all members of the Kakī Recovery Programme and the Orange-fronted Parakeet Recovery Programme for their assistance with sample collection and their ongoing support. This research has been funded by the Ministry of Business, Innovation and Employment (MBIE) Endeavour Fund (UOCX1602 awarded to TES), the Brian Mason Scientific and Technical Trust (awarded to SJG and TES), and the Mohua Charitable Trust (awarded to TES).

## ACCESSIBILITY

Genomic data provided in this manuscript are available through a password protected server on the Conservation, Systematics and Evolution Research Team’s website (http://www.ucconsert.org/data/). Kakī and kākāriki karaka are taonga (treasured) species. For Māori (the indigenous people of Aotearoa), all genomic data obtained from taonga species have whakapapa (genealogy that includes people, plants and animals, mountains, rivers and winds) and are therefore taonga in their own right. Thus, these data are tapu (sacred) and tikanga (customary practices, protocols, and ethics) determine how people interact with it. To this end, the passwords for the genomic data in this manuscript will be made available to researchers on the recommendation of the kaitiaki (guardians) for the iwi (tribes) that affiliate with kakī and kākāriki karaka.

## AUTHOR CONTRIBUTIONS

This research was led by SJG, AWS, and TES, with the research concept developed in collaboration with all authors. All DNA extractions and microsatellite lab work was completed by SJG. The kakī pedigree was developed by SJG in collaboration with LB and SC. The kākāriki karaka pedigree was developed by AR with edits by SJG. The kākāriki karaka genome was assembled by RM. SJG and RM conducted SNP discovery. All statistical analyses were conducted by SJG. The manuscript was written by SJG in collaboration with all authors.

