## Supplemental Materials for "A comparison of pedigree, genetic, and genomic estimates of relatedness for informing pairing decisions in two critically endangered birds"

### S1.1: COANCESTRY Microsatellite Simulations

The programme COANCESTRY v. 1.0.1.9 (Wang et al. 2011) offers seven different estimators of relatedness for genetic and genomic markers, and to choose the most appropriate estimator for the kakī and kākāriki karaka microsatellite datasets, we employed the simulation module within COANCESTRY using allele frequencies, missing data, and error rates from our microsatellite datasets. To produce dyads that represent the relationships and degree of inbreeding found within kakī and kākāriki karaka, we used the R package 'identity' (Li 2010) to generate 10,879 dyads for kakī and 1,484 dyads for kākāriki karaka based on the pedigrees of both species. The frequency of each unique dyad in the kakī and kākāriki karaka datasets were scaled to create 1,000 dyads for each set that are representative of relationships between individuals used in this study. The COANCESTRY simulations were conducted using allele frequencies, error rates, and missing data rates from each microsatellite data set, with settings changed to account for inbreeding. The triadic likelihood approach (Wang 2007) was selected given it had the highest Pearson's correlation with 'true' relatedness and lowest variance for both kakī (Table S1) and kākāriki karaka (Table S2), as per Hammerly et al. (2013).

Table S1. Average relatedness, variance, and correlation to 'true' relatedness for seven relatedness estimators using simulated datasets from kakī microsatellite allele frequencies and dyads in the kakī pedigree.

| | TrioEst | WEst | LLEst | LREst | REst | QGEst | MEst | Actual Relatedness |
|---|---|---|---|---|---|---|---|---|
| Average | 0.214 | 0.063 | 0.064 | 0.067 | 0.065 | 0.064 | 0.236 | 0.07 |
| Variance | **0.047** | 0.131 | 0.125 | 0.088 | 0.098 | 0.103 | 0.051 | 0.023 |
| Pearson's R | **0.511** | 0.407 | 0.418 | 0.483 | 0.473 | 0.451 | 0.502 | 1 |

Table S2. Average relatedness, variance, and correlation to 'true' relatedness for seven relatedness estimators using simulated datasets from kākāriki karaka microsatellite allele frequencies and dyads in the kākāriki karaka pedigree.

| | TrioEst | WEst | LLEst | LREst | REst | QGEst | MEst | Actual Relatedness |
|---|---|---|---|---|---|---|---|---|
| Average | 0.283 | 0.145 | 0.149 | 0.159 | 0.15 | 0.152 | 0.315 | 0.151 |
| Variance | **0.07** | 0.168 | 0.166 | 0.123 | 0.158 | 0.146 | 0.076 | 0.051 |
| Pearson's R | **0.675** | 0.567 | 0.568 | 0.663 | 0.533 | 0.597 | 0.661 | 1 |

### S1.2: SNP Filtering

Base filtering was applied to all tried filtering datasets, including a filtering to retain only biallelic SNPs with a minor allele frequency (MAF) greater than 0.05, a quality score greater than 20, and maximum missingness of 10% per site. This base filtering was designed to increase the quality, completeness, and reliability of SNPs by removing low quality markers and potential artefacts captured by rare multiallelic sites (Campbell et al. 2016) or low frequency sequencing error.

Different depth filters for each site were tested for each dataset to achieve an average of ~10x depth across all sites for each individual, to provide more certainty around reference-guided SNPs used here. Preliminary testing with kakī revealed that using an average minimum depth of 10x was sufficient for meeting this criteria (e.g., the lowest per individual average depth observed post filtering was 9.6x, with mean depth across all sites and individuals of 28.7x ± 10.29 SD). This depth was also used for an aligned study (see Galla et al. 2019). Applying this same filtering scheme to the kākāriki karaka dataset did not achieve an average depth of ~10x across all sites for each individual (the minimum average depth = 5.44x, mean depth across all sites and individuals = 15.93x ± 10.11). Therefore, a filtering trial using two different depth filtering strategies was employed, with one filtering using a hard minimum cut-off of 5x depth and the other using an average minimum depth of 20x. Both of these filtering trials also employed a hard maximum cutoff of 200x depth, to filter obvious high-coverage sites that come from collapsed repeats in the reference genome. Because there are known parent-offspring relationships represented in our dataset, and parent-offspring genomic contribution is 50%, we used relatedness between parents and offspring as a biologically meaningful measure to understand which approach best approximated 0.5 with the greatest precision. It was found that a hard minimum cut-off of 5x per SNP resulted in relatedness estimates that were more accurate and precise (Table S3).

Table S3: Results from filtering trials for depth, $r^2$, and HWE using the kākāriki karaka data set. Base filtering refers to filter steps for biallelic SNPs with a MAF of 0.05, and missingness of 0.1. Scaled refers to the chosen data set, after scaling for self-relatedness to be equal to 1. Bold text refers to filtering scheme chosen. Alternating blue and white cell fill denotes trials with aligned depth and $r^2$ settings.

| Trial | Average *R* ± SD | Average PO *R* ± SD | Min PO *R* | Max PO *R* |
|---|---|---|---|---|
| Base filtering, minimum depth of 5x, maximum depth 200x, r2 0.4 | 0.25 ± 0.1 | 0.43 ± 0.04 | 0.36 | 0.57 |
| Base filtering, minimum depth of 5x, maximum depth 200x, r2 0.4, HWE | 0.02 ± 0.22 | 0.43 ± 0.11 | 0.19 | 0.67 |
| **Base filtering, minimum depth of 5x, maximum depth 200x, r2 0.6** | **0.24 ± 0.10** | **0.43 ± 0.04** | **0.35** | **0.58** |
| **Base filtering, minimum depth of 5x, maximum depth 200x, r2 0.6, Scaled** | **0.29 ± 0.12** | **0.53±0.03** | **0.47** | **0.67** |
| **Base filtering, minimum depth of 5x, maximum depth 200x, r2 0.6, HWE** | **0.02 ± 0.22** | **0.43 ± 0.12** | **0.19** | **0.68** |
| Base filtering, minimum depth of 5x, maximum depth 200x, r2 0.8 | 0.24 ± 0.11 | 0.44 ± 0.04 | 0.36 | 0.59 |
| Base filtering, minimum depth of 5x, maximum depth 200x, r2 0.8, HWE | 0.02 ± 0.23 | 0.45 ± 0.12 | 0.21 | 0.7 |
| Base filtering, average minimum depth 20x, maxiumum depth 200x $r^2$ 0.4 | 0.08 ± 0.13 | 0.33 ± 0.08 | 0.17 | 0.53 |
| Base filtering, average minimum depth 20x, maxiumum depth 200x $r^2$ 0.4, HWE | 0.01 ± 0.19 | 0.36 ± 0.11 | 0.12 | 0.58 |
| Base filtering, average minimum depth 20x, maxiumum depth 200x $r^2$ 0.6 | 0.05 ± 0.15 | 0.33 ± 0.09 | 0.14 | 0.55 |
| Base filtering, average minimum depth 20x, maxiumum depth 200x $r^2$ 0.6, HWE | 0.01 ± 0.19 | 0.36 ± 0.12 | 0.12 | 0.59 |
| Base filtering, average minimum depth 20x, maxiumum depth 200x $r^2$ 0.8 | 0.03 ± 0.17 | 0.34 ± 0.07 | 0.21 | 0.57 |
| Base filtering, average minimum depth 20x, maxiumum depth 200x $r^2$ 0.8, HWE | 0.00 ± 0.20 | 0.37 ± 0.12 | 0.12 | 0.62 |

In addition to depth for kākāriki karaka, different $r^2$ filters (i.e., $r^2$ = 0.4, 0.6, and 0.8) for linkage disequilibrium were applied to see how this variable affected relatedness estimates. Further, HWE filters of 0.05 using KGD were applied to each of these trials to see if using a HWE filter on each of these approaches affected the accuracy and precision of relatedness (Tables S3 and S4). Overall, using a HWE filter resulted in less accurate and precise estimates of relatedness, which may be attributed to our datasets consisting mostly of family groups, thereby violating the assumption of random breeding. Strong to moderate LD filters ($r^2$ = 0.4, 0.6) produced more accurate and precise estimates of relatedness in kakī, but did not make a substantial difference in kākāriki karaka. A moderate LD filter ($r^2$ = 0.6) was chosen for both datasets.

Table S4: Results from filtering trials for $r^2$ and HWE using the kakī data set. Base filtering refers to set filters for biallelic SNPs with a MAF of 0.05, an average mean depth of 10x, and missingness of 0.1. Scaled refers to the chosen data set, after scaling for self-relatedness to be equal to 1.

| Trial | Average *R* ± SD | Average PO *R* ± SD | Min PO *R* | Max PO *R* |
|---|---|---|---|---|
| Base filter, $r^2$ 0.4 | 0.33 ± 0.07 | 0.52 ± 0.03 | 0.47 | 0.60 |
| Base filter, $r^2$ 0.4, HWE | 0.02 ± 0.12 | 0.37 ± 0.06 | 0.30 | 0.55 |
| **Base filter, $r^2$ 0.6** | **0.25 ± 0.08** | **0.47 ± 0.04** | **0.42** | **0.59** |
| **Base filter, $r^2$ 0.6, Scaled** | **0.27 ± 0.09** | **0.54 ± 0.03** | **0.49** | **0.61** |
| Base filter, $r^2$ 0.6, HWE | 0.01 ± 0.13 | 0.38 ± 0.08 | 0.28 | 0.59 |
| Base filter, $r^2$ 0.8 | 0.16 ± 0.09 | 0.42 ± 0.06 | 0.34 | 0.58 |
| Base filter, $r^2$ 0.8, HWE | 0.00 ± 0.13 | 0.37 ± 0.09 | 0.26 | 0.60 |

### S1.3: SNP-based relatedness estimates

To produce pairwise estimates of relatedness using whole-genome SNPs, we used the R script KGD (Dodds et al. 2015), as it was designed to estimate relatedness using reduced-representation and resequence data while taking into account read depth. In order to evaluate the performance of KGD, estimates of relatedness were produced and compared with the triadic likelihood approach method (i.e., TrioML; Wang 2007), as it is one of the few estimators that accounts for instances of inbreeding. The R package *related* (Pew et al. 2015) was used instead of the programme COANCESTRY (Wang et al. 2011) to produce TrioML estimates using genome-wide SNPs, as it is able to handle more than 20,000 markers while still using the same base-code as COANCESTRY. Settings were set to account for inbreeding and calculate 95% confidence intervals with a bootstrap value of 100. Known parent/offspring dyads were used to evaluate precision, as parents/offspring relatedness should approximate 0.5 (Speed & Balding 2015).

Results indicate that KGD was able to produce estimates of parent-offspring relatedness that more closely approximated 0.5 than TrioML, with a smaller standard deviation and range of parent/offspring relatedness values produced (Table S3). While KGD estimates were more precise for parent-offspring relationships than TrioML, these estimates still significantly correlate with the TrioML estimator for both kakī (Pearson's $r$ = 0.96, $p$ < 0.001) and kākāriki karaka (Pearson's $r$ = 0.96, $p$ < 0.001).

Table S5: SNP-based estimates of relatedness using KGD and TrioML approaches in kakī and kākāriki karaka.

| Species | Estimator Used | Average *R* | Min. *R* | Max. *R* | Average Parent - Offspring *R* | Min. Parent - Offspring *R* | Max. Parent - Offspring *R* | Pearson's Correlation *r* |
|---|---|---|---|---|---|---|---|---|
| Kakī | KGD | 0.27 ± 0.09 | 0.127 | 0.612 | 0.541 ± 0.033 | 0.489 | 0.612 | 0.96 |
| | TrioML | 0.06 ± 0.06 | 0 | 0.403 | 0.265 ± 0.058 | 0.168 | 0.403 | |
| Kākāriki karaka | KGD | 0.29 ± 0.12 | 0.08 | 0.67 | 0.53 ± 0.03 | 0.47 | 0.67 | 0.96 |
| | TrioML | 0.07 ± 0.10 | 0 | 0.43 | 0.26 ± 0.04 | 0.18 | 0.43 | |

### S1.4: MSI Correlations

Correlations between pedigree-, microsatellite-, and SNP-based MSI scores were performed in the manuscript. Scatterplots showing these relationships are provided below.

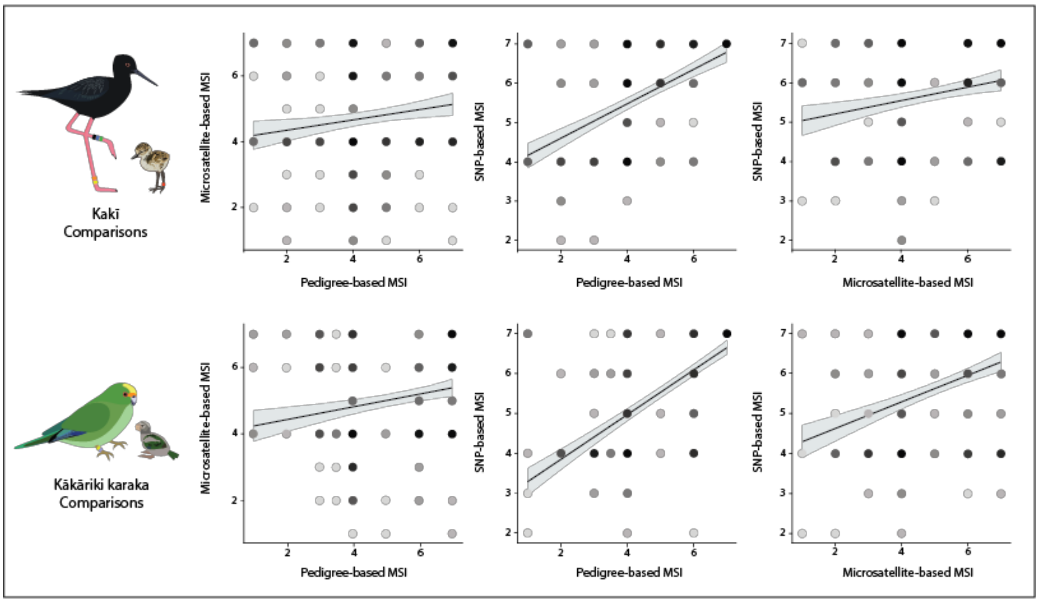

*Figure S 1. Scatterplots showing relationships between pedigree-, microsatellite-, and SNP-based MSI scores in kakī  and kākāriki karaka. Darker points denote higher frequencies than lighter points. A trend line (black) and 95% confidence intervals (grey) are shown in each comparison.*
